# Atomic force microscopy reveals the role of vascular smooth muscle cell elasticity in hypertension

**DOI:** 10.1101/2020.12.01.407254

**Authors:** Yi Zhu

## Abstract

The vascular smooth muscle cell (VSMC) mechanical properties not only provide intrinsic cellular functions, but also influence many vascular and circulation functions in physiology. In this report, the VSMCs of thoracic aorta from 16 week age Wistar-Kyoto normotensive rats (WKY) and spontaneously hypertensive rats (SHR) were used as research subjects to reveal hypertension mechanism at a single cell level using atomic force microscopy (AFM). The apparent elastic modulus was significantly increased in VSMCs from SHRs compared to those from WKYs. Treatment with cytochalasin D (CD), ML7, Y27632 and lysophosphatidic acid (LPA) modulated VSMC stiffness of WKYs and SHRs. A spectral analysis approach was applied to further investigate the time-dependent change in VSMC elasticity of WKYs and SHRs. This report demonstrated the efficacy of real-time analysis of VSMC elasticity by AFM nano-indentation, and revealed real-time functional differences in biomechanical characteristics of VSMCs with drug treatments.

## 1. Introduction

Vascular smooth muscle cells (VSMCs) locate blood vessel medial layer as a main component and bear mechanical stress and pressure from blood flow, and sustain vascular tone and resistance. A number of recent studies have demonstrated that changes in a cell’s elastic characteristics can affect its response to the external mechanical force (Hill M.A., et al., 2016; Dhar S., et al., 2017). The single-cell mechanical property and behavior of VSMC is chiefly considered to play a crucial role in the development of vascular diseases, and atomic force microscopy (AFM) is currently the most wonderful tools for determining this interaction (Zhu W., et al., 2018; Leloup A.J.A., et al., 2019; Sanyour H.J., et al., 2020).

The VSMC intrinsic properties not only perform a normal cellular function to sustain and support vascular geometric architecture, but also take some important actions to participate the regulation of biophysical and biochemical properties for blood vessel (Jia G., et al., 2015; Zhang J., et al., 2016; Yang J., et al., 2017). With the development and applications of AFM technology, people gradually concentrate their research on reconstituted tissues and single cell detections (Huang H., et al., 2018; Li N., et al., 2018; Zhou Z., et al., 2020). Hypertension is a common age-related vascular disease, and many factors can induce age-related vascular dysfunctions and diseases (Touyz R.M., et al., 2018). However, the detailed mechanisms that induce hypertension still need to be elucidated. Currently, people attempt to analyze and reveal the hypertension mechanism in single molecule and single cell level (Huang H., et al., 2018; Zhu Y., et al., 2018 and 2019). The cytoskeleton contents, the polymerization and arrangement of actin filaments were directly responsible for the VSMC elasticity (Shen K., et al., 2019; Rickel A.P., et al., 2020). The investigation in single cell level can supplement studies on complicated living organisms or an intact tissue to determine the related pathways that regulate cell elasticity and adhesion (Zhou N., et al., 2017 a and b). Furthermore, cells are in micro-scales and easy to break, and AFM provides a probability to manipulate VSMC at an individual cell level due to its nano-sensitivity under liquid environment (Sanyour H., et al., 2018, 2019 and 2020). The experimental medicines are administered in micro-volume by a pipette and ensured drugs to diffuse and aim the measured cells, and people can fully and perfectly employ AFM to perform a continuous real-time measurement in the absence and presence of drugs on a single cell. The single spectral analysis is an approach to reveal mathematical decomposition of the elasticity waveform and further demonstrates the underlying molecular mechanism (Hong Z., et al., 2015; Sehgel N.L., et al., 2015a and b). The drug cytochalasin D (CD) depolymerizes and breaks apart actin filaments, and the drug ML7 dephosphorylates myosin light chain to inhibit the establishment of actin binding with myosin (Zhang J., et al., 2016). Additionally, the drug Y27632 inhibits the Rho-associated protein kinase (ROCK) and the drug lysophosphatidic acid (LPA) enhances integrin proteins to adhere the extracellular matrix (ECM) and activates Rho kinase to phosphorylate myosin light chain kinase (MLCK) (Staiculescu M.C., et al., 2014; Turner C.J., 2015). In this report we chose these drugs using AFM to measure the stiffness of thoracic aortic VSMCs *in vitro*. An investigation was taken to study and reveal the real-time record of single VSMC mechanical property and behavior. Moreover, we analyzed and interpreted the oscillatory waveforms of VSMC elasticity for various drug treatments to reveal the underlying cellular and molecular mechanisms of VSMC stiffness in hypertension by a spectral analysis approach.

## 2. Materials and Methods

### 2.1 Vascular smooth muscle cell isolation, cell culture, and treatments

Male WKYs and SHRs at 16–18 weeks of age were utilized in this study. All animal procedures were done under the *Guide for the Care and Use of Laboratory Animals* (NIH 85-23, revised 2011). Primary VSMCs from thoracic aorta of three experimental WKY and SHR rats were enzymatically isolated and cultured in Dulbecco’s Modified Eagle’s medium (DMEM) with 10% fetal bovine serum (FBS), 10 mmol/L HEPES, 2 mmol/L L-glutamine, 1 mmol/L sodium pyruvate, 100 U/mL penicillin, 100 μg/mL streptomycin, and 0.25 μg/mL amphotericin B and used at passages 2 to 4 (Sanyour H.J., et al., 2020).

### 2.2 VSMC image and stiffness measured by AFM

Single VSMC image and lively measurements of cell elasticity were operated in contact mode by an AFM instrument, which is a Bioscope System (Model IVa, Veeco Mertrology Inc., Santa Barbara, CA) mounted on an Olympus IX81 microscope (Olympus Inc., NY). The employed AFM probes were silicon nitride microlevers (Model micro lever cantilever, Veeco Mertrology Inc., Santa Barbara, CA; spring constant ranging 10-30 pN/nm) and purchased from Veeco Mertrology Inc. (Santa Barbara, CA). The AFM tip was put in the mid-site between VSMC margin and the nucleus for nano-indentation to measure WKY and SHR elasticity. The AFM probe was continuously indented 2 minute to collect force curves for determining the mean stiffness of individual WKY and SHR VSMC, and the experimental VSMCs from three rats were assessed then averaged together for the stiffness of WKYs and SHRs. The force curves were interpreted using proprietary software NForceR (registration number TXu1-328-659), and the VSMC elastic modulus was translated from these force curves into Young’s modulus using a modified Hertz model. The calculation of the elastic modulus was:

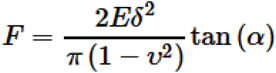

where the indentation force (F) was stated and described using Hooke’s law (F =κΔx, κ and Δx denote the AFM probe’s spring constant and the probe’s apparent deflection). The indentation depth (δ) is identified from the difference in the AFM piezo movement in z direction and the AFM probe deflection. E is the Young’s modulus of experimental cell as the value of elasticity, and ν denotes 0.5 for cell as the Poisson ratio. The numerical α is the semi-included angle of the cone for a pyramidal tipped probe and determined by the probe shape.

### 2.3 Dynamic stiffness in single VSMC measurement by AFM

The experimental VSMCs were nano-indented for the duration of 30 minute to examine the temporal characteristics of the cell stiffness, and then VSMCs were treated in micro-volume by a pipette with CD (10 μmol/L; Sigma, St. Louis, MO), with ML7 (10 μmol/L; Sigma, St.

Louis, MO), with Y27632 (5μmol/L; Sigma, St. Louis, MO), with LPA (2 μmol/L; Sigma, St. Louis, MO) for another 30 minute continuous AFM investigation. The curves were continuously recorded and collected during the whole measuring procedure, and applied to determine elastic stiffness, absence and presence of drugs. A spectral analysis procedure was exploited for analysis and following translation of the oscillation waveforms for elasticity data, and linear trends were evaluated and subtracted from each series ahead of a spectral analysis. To reveal the average group behavior of the oscillations, three values of amplitude, frequency and phase for every experimental subject were further investigated and averaged: phases (φ)as a simple mean; frequencies (f) were converted to periods (1/f) ahead of averaging; amplitudes (A) were log10-transformed before averaging the mean. The mean period and mean log-amplitude were then transformed back to frequency and amplitude. A composite time series for each treatment set was constructed as:

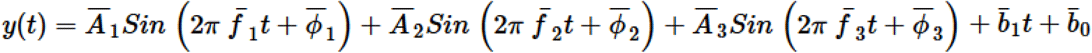

where *b*_1_ and *b*_0_ denote respectively the slope and intercept of the linear trend, and the bar above each component indicates the average value (Zhu Y., et al., 2018). A brief explanation for the singular spectrum analysis equation and application was provided in this report, and we detail stated and described the measurement of dynamic stiffness by AFM in single VSMC and the analysis of oscillation waveform by singular spectrum analysis.

### 2.4 Statistical analysis

Data are expressed as mean ± SEM for the number of samples reported in this report. Statistically significant differences between WKYs and SHRs were determined by Student’s t-test. A value of P<0.05 was considered a significant difference.

## 3. Results and Discussion

### 3.1 VSMC AFM image and topography

The SHR developed from WKY rat as an animal model for specific studies of cardiovascular disease, thus we analyzed and compared heights and topographic images of VSMCs isolated from thoracic aorta of WKYs (n=4, from 3 rats) and SHRs (n=5, from 3 rats). The VSMC surface areas of WKYs vs. SHRs were 10695±339 μm2 vs. 12380±483 μm2, and the VSMC surface area of SHRs was significantly larger than WKYs (p<0.05). For the height measurement, we set AFM to predetermine the line across the cell and take a 30 second period reading. Waited 600 seconds (10 minutes), and started recording line scans of height again for another 30-second period. We repeated this procedure for many times to obtain the VSMC height (Figure 1). The VSMC topography and shape of SHRs showed to be larger and higher than WKYs (p<0.05) due to α-SMA over production and F-actin over assembly. The expression of cytoskeletal actin in SHRs is obviously higher than (p<0.05) WKYs at the same age and the denser actin filaments make a lot of crosslinking polymers inside VSMCs (Sehgel N. L., et al., 2013).

**Figure 1.**
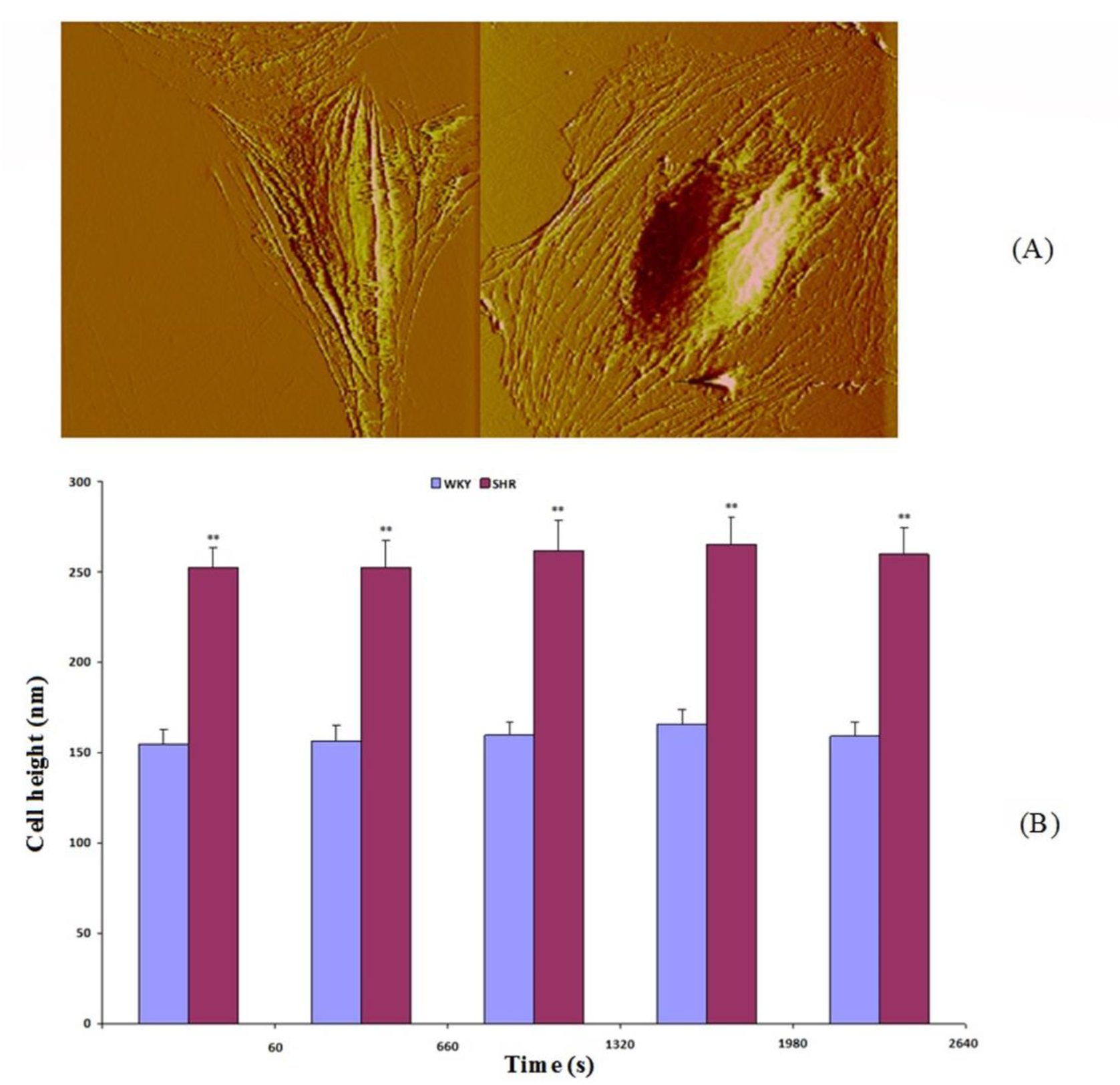
Topographic characterizations of WKY and SHR VSMCs. (A) Example of WKY TA (left) and SHR TA VSMC (right) AFM deflection image. (B) Scanning height data of WKY TA (n=5, from 3 rats) and SHR TA (n=5, from 3 rats). **p<0.01 (SHR TA VSMC compared to WKY TA VSMC in different time regions).

### 3.2 Drugs effect on VSMC elasticity

The VSMC elasticity data are consistent with the above topographical observations, and indicate that the intrinsic property of a single cell reflects its mechanical characteristics. By CD and ML7 evaluations, the elasticities of VSMCs were dramatically reduced and there were no significant differences between WKYs and SHRs (Sehgel N. L., et al., 2013). The drug Y27632 (5μmol/L) was performed to treat WKY and SHR VSMCs, VSMC elasticity of SHRs showed a higher value in presence of Y27632 in comparison to that of WKYs (p<0.005, Figure 2A), whereas the drug lysophosphatidic acid (LPA) (2 μmol/L) increased VSMC elasticity in both WKYs and SHRs, but to a larger extent in SHR (p<0.001, Figure 2A).

**Figure 2.**
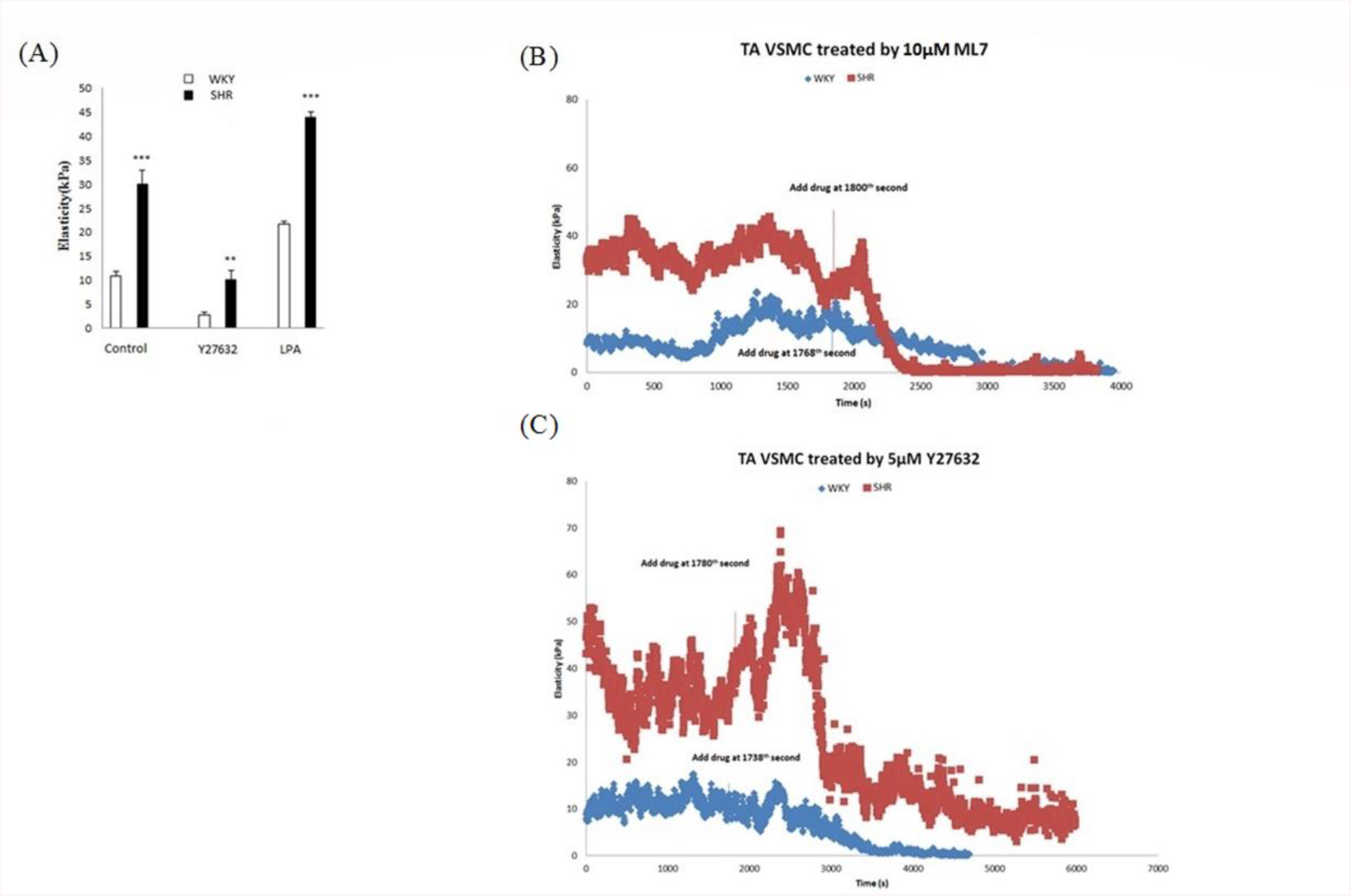
(A) VSMCs were treated with 5μmol/L Y27632 to inhibit the ROCK, with 2 μmol/L LPA to increase integrin adhesion to ECM for 30 minute measurement. **P<0.005 (WKY vs. SHR), ***P<0.001(WKY vs. SHR). (B) Examples of real-time cell elastic modulus for typical WKY (blue) and SHR (red) vascular smooth muscle cells using 10μmol/L ML7 treatment and (C) 5μmol/L Y27632 treatment are shown.

The time series behavior in VSMC elasticity of WKYs and SHRs with 10 μmol/L CD, 10 μmol/L ML7 (Figure 2B), 5μmol/L Y27632 (Figure 2C) and 2 μmol/L LPA treatments in single cell level was further investigated by a spectral analysis approach. After 10 μmol/L CD treatment there were not any significant differences in three components of VSMC elasticity oscillatory behaviors between WKYs and SHRs. Interestingly, in the second component the amplitude of WKY was higher than that of SHR (p<0.05) by CD treatment (Figure 3A). Possibly CD depolymerizes and disrupts actin filaments in SHR cells, and shows a lower amplitude in the second component. The mechanism will be further revealed. Additionally, ML7 is a drug to inhibit myosin light chain phosphorylation, and is applied to VSMC to test its effect on SHR and WKY VSMC elasticity. After 10 μmol/L ML7 treatment, there were also not any significant differences between WKY and SHR VSMCs in three components of oscillatory behaviors. In three principle components, the frequencies and amplitudes of both WKYs and SHRs were not significantly different (p>0.05) (Figure 3B).

**Figure 3.**
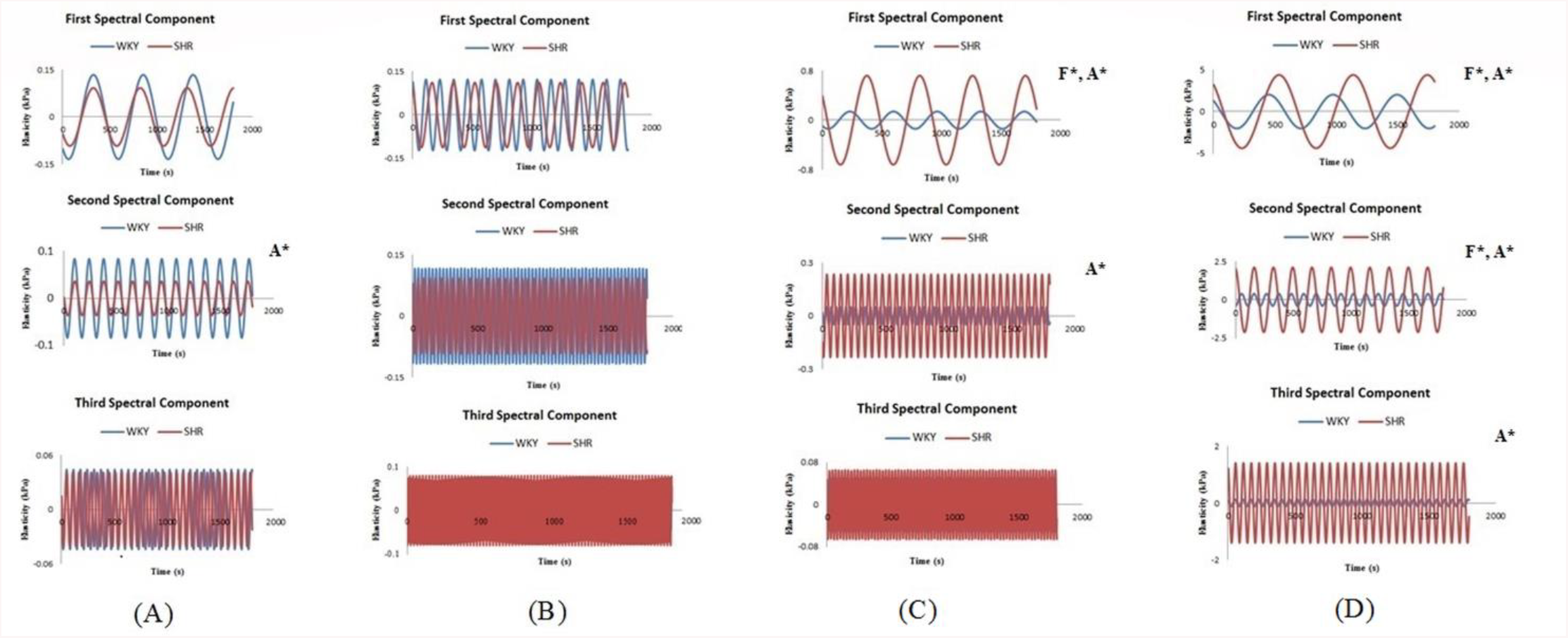
Over a 30 minute period, mathematical analysis of the elastic modulus waveform in the presence of drugs indicated three principle components of oscillation by spectral analysis for WKY and SHR thoracic aorta vascular smooth muscle cells. (A) WKYs (5 cells from 3 animals) and SHRs (4 cells from 3 animals) treated by 10 μM CD. (B) WKYs (9 cells from 3 animals) and SHRs (5 cells from 3 animals) treated by 10 μMML7. (C) WKYs (5 cells from 3 animals) and SHRs (6 cells from 3 animals) treated by 5μM Y27632. (D) WKYs (5 cells from 3 animals) and SHRs (5 cells from 3 animals) treated by 2 μM LPA. F* and A* indicate that the frequency and amplitude of the component were significantly different from WKYs and SHRs in the presence of drugs, *P* < 0.05.

From an individual cell point of view, SHRs strongly and vehemently responded to both LPA and Y27632 treatments. The spectral analysis clearly demonstrated the dynamic oscillatory behaviors in VSMC elasticity that are driven by actin–ECM interactions at the absence and presence of these two drugs. The drugs activate or inactivate VSMC elastic characters through a series of cascade responses, thus after drug treatments the frequencies and amplitudes of first spectral component (large visible oscillation, Figure 3C and D) between SHRs and WKYs existed significant differences (p<0.05). Moreover, the amplitudes of the second spectral component between SHRs and WKYs existed significant differences (p<0.05) (Figure 3C and 3D).

Rho kinase acts as a signaling molecule to influence VSMC stiffness via the Ca2+-CaM-MLCK pathway and with some cascade cycles by phosphorylation and dephosphorylation. Rho kinase also regulates aortic VSMC stiffness via actin/ serum response factor (SRF)/myocardin in hypertension (Zhou N., et al., 2017b). The WKYs and SHRs are at 16-18 weeks of age, from the prior reports the basal expression level of myosin light chain (MLC) in both WKYs and SHRs was closed, but the expression of pMLC was found to be increased in SHRs (Sehgel N. L., et al., 2013). The drug Y27632 inhibits Rho kinase to phosphorylate MLCK and dephosphorylates pMLC, indirectly keeps actin away from binding with myosin to depolymerize the establishment of actin-myosin complex and eliminates the VSMC elasticity (Pierce G.L. 2017). In addition, the ECM-integrin-cytoskeletal axis is an important pathway to influence VSMC stiffness by regulating α-SMA expression, and the coordinate ability of ECM-integrin-actin is attenuated to produce hypertension (Zhu Y., et al., 2018 and 2019). The cytoskeletal α-smooth muscle actin (α-SMA) importantly responds mechanical forces through ECM-integrin-cytoskeletal axis to mediate VSMC stiffness, and it is over expressed to be a decisive factor leading to an increase in aortic stiffness for inducing hypertension. The α-SMA expression and polymerization in SHR VSMCs are obviously higher than WKYs at the same age (Sehgel N. L., et al., 2013).

Attenuating the stiffness of VSMC via several pathways, Y27632 gently damaged the crosslinking of α-SMA filaments and the contractility of myosin production in comparison to the drug CD and ML7, consequently VSMC elasticity of SHRs showed a higher value in presence of Y27632 in comparison to that of WKYs.

The drug lysophosphatidic acid (LPA) activates Rho kinase to phosphorylate MLCK and promote α-SMA polymerization (Staiculescu M.C., et al., 2014). Meanwhile, LPA enhances integrin to adhere to the ECM and activate Rho kinase. Increasing adhesion interaction between α5β1integrin and fibronectin (one component of ECM) is related to increasing cell stiffness via ECM-integrin-cytoskeletal axis, and MLCK was also found to be over expressed in VSMCs of SHRs (Hong Z.K., et al., 2012 and 2013; Sehgel N. L., et al., 2013 and 2015a). LPA increases α5β1integrin to adhere to ECM for promoting actin expression and polymerization. The interaction between α5β1 integrin and FN is specific and important in the mechanical transduction of VSMC, and α5β1 integrin is the major receptor for FN (Sun Z., et al., 2005; Wu X., et al., 1998 and 2001). The α5β1integrin provides the bio-mechanical linkage between α-SMA and fibronectin (FN) in extracellular space, and α-SMA responds the bio-mechanical forces through integrin-mediated cell-ECM interactions to alternate cytoskeleton system of VSMCs (Hartman C.D., et al., 2016; Hays T.T., et al., 2018). The ECM-integrin-cytoskeletal axis and the contractility of myosin production are two independent pathways to regulate the VSMC stiffness (Huang H., et al., 2018). The α-SMA was highly expressed in the VSMC of SHRs, moreover, MLCK was also found to be over expressed in SHRs to stiffen the VSMC due to enhancing the VSMC contractile process (Rodenbeck S.D., et al., 2017), thus this analysis showed VSMC elastic moduli of SHRs were greater than (p<0.05) those of WKYs.

Previous studies have shown that both internal and external biomechanical forces can act through the cytoskeleton, thereby affecting local elasticity and cell behavior (Fletcher D.A., et al., 2010). The differences in stiffness and time-dependent oscillations were largely influenced by actin cytoskeletal dynamics. Dynamical alternation of α-SMA constructs different high-level linkage structures in VSMCs, affecting cell elasticity and cellular stress relaxation behavior (Sun Z., et al., 2008 and 2012; Wu X., et al., 2010). To further verify the internal characteristics of cells that reflect the mechanical properties of cells, various drug treatments that affect the cytoskeleton and corresponding vascular smooth muscle contraction mechanisms have been carried out on VSMCs for in-depth research. Three spectral components are determined in the oscillation mode, so it is reasonable to assume that more than one mechanism causes the spontaneous oscillation of cell elasticity, and therefore may play a role in the increased vascular stiffness observed in hypertension. The elastic oscillations of VSMCs represent the inherent characteristics of cells and involve the cytoskeleton structure responsible for the interaction of actin-ECM and actin-myosin. At the same time, the oscillation of VSMC elasticity reveals the polymerization and depolymerization of α-SMA. Different pharmacological mechanisms produced the different individual cell elastic behavior after the drug treatments. Spectral analysis showed that compared with WKY rats, SHRs usually have lower frequencies and larger amplitudes. The general pattern of slow, larger oscillations in SHRs and faster, smaller oscillations in WKY VSMCs (Sehgel N. L., et al., 2013). After Y27632 treatment, the frequency of the first wave component is significantly reduced in SHR VSMCs, whereas the amplitudes of the first and second wave components are increased in SHR VSMCs. The frequency and amplitude showed not to be a significant difference between WKY and SHR VSMCs in the third wave component. All in all, the spectral analysis indicated that Y27632 gently attenuated VSMC stiffness. The drug LPA polymerizes α-SMA to increase VSMC stiffness, from the spectral analysis the amplitudes of all three wave components are enhanced in SHR VSMCs, and the frequencies of the first and second wave components are significantly reduced in comparison to WKY VSMCs. The drug CD breaks apart the actin cytoskeletal network and the drug ML7 dephosphorylates myosin light chain to block the interaction between actin and myosin. These two drugs CD and ML7 strongly and irreversibly destroy the cross-linking of actin filaments and the contractility produced by myosin. Both WKYs and SHRs completely lose their elasticity, and therefore exhibit inactivity in the three components of the oscillation.

In summary, cellular mechanisms underlying differences in VSMC stiffness were investigated using AFM. For decades, pharmacists have developed many drugs for the treatment of certain vascular diseases based on the role of the actin-integrin axis in the mechanical properties of VSMCs, and AFM provides a way to manipulate an individual VSMC due to its nano-sensitivity under physiological condition and liquid environment. At present, people have used AFM in many applications to study the mechanical properties of a single VSMC, and the intrinsic changes of VSMC enable people to open up new therapeutic ways for the treatment of multiple diseases and update our understanding of vascular biology (Nance M.E., et al., 2015; Pierce G.L. 2017). The future trends of employing AFM tip coating techniques for adhesive assessment and super-resolution fluorescence microscopy for cytoskeletal tracking will further resolve individual VSMC elasticity and its role in physiological process of living organisms (Ella S.R., et al., 2010). The *in vivo* mechanism of how individual VSMC elasticity to regulate vascular processes is still unknown, thus the AFM detection *in vitro* combined with the investigation *in vivo* by other techniques will also provide a perspective view to describe the VSMC elastic characters in the coming researches (Lacolley P., et al., 2017 and 2018).

## LIST OF ABBREVIATIONS

AFM: atomic force microscopy
α-SMA: α-smooth muscle actin
CD: Cytochalasin D
DMEM: Dulbecco’s Modified Eagle’s medium
ECM: extracellular matrix
FBS: fetal bovine serum
HEPES: 4-(2-hydroxyethyl)-1-piperazineethanesulfonic acid
LPA: lysophosphatidic acid
ML7: Hexahydro-1-[(5-iodo-1-naphthalenyl) sulfonyl]-1*H*-1, 4-diazepine
MLCK: myosin light chain kinase
ROCK: Rho-associated protein kinase
SHR: spontaneously hypertensive rat
VSMC: vascular smooth muscle cell
WKY: wistar-kyoto normotensive rat
Y27632: (1R, 4r)-4-((R)-1-aminoethyl)-N-(pyridin-4-yl) cyclohexanecarboxamide

